# Whole exome-sequencing of vitiligo lesions indicate lower burden of somatic variations: implications in risk for non-melanoma skin cancers

**DOI:** 10.1101/2022.05.28.493819

**Authors:** Iti Gupta, Shambhavi Shankrit, Kiran Narta, Madeeha Ghazi, Ritika Grover, Rajesh Pandey, Hemanta K Kar, Shruti M Menon, Aayush Gupta, Vamsi K Yenamandra, Archana Singh, Mitali Mukerji, Arijit Mukhopadhyay, Rajni Rani, Rajesh S Gokhale, Debasis Dash, Vivek T Natarajan

## Abstract

Mapping of somatic variations has enabled understanding the progression of clonal variations from healthy skin to cutaneous malignancies. Highlighting, the adaptive nature of pigmentation, germline mutations in albinism amplify skin cancer susceptibility. However, lower incidence of non-melanoma skin cancer among subjects with acquired depigmenting skin disorder vitiligo is enigmatic and a matter of longstanding debate. To address this, we performed high-coverage exome sequencing of matched non-lesional and lesional vitiligo skin along with whole blood to account for germline variations. Our analysis suggests lower burden of somatic cancer-associated variations in exposed depigmented lesional skin compared to the non-lesional skin. A detailed investigation of vitiligo skin transcriptome reveals elevation of DNA repair and cell-proliferation pathways. Validation by comet-assay for DNA damage and cell cycle analysis of epidermal cells suggest undamaged DNA in vitiligo lesions that could be attributed to higher proliferation-coupled repair. Endorsing this, UV-signature variations are not prominent, instead SBS5 associated with endogenous mutational processes is conspicuous in both the vitiligo tissues. Our systematic pilot study indicates lower somatic mutation burden in vitiligo skin and supports the earlier demographic observation on lower risk of non-melanoma skin cancer in vitiligo subjects, providing an opportunity to learn strategies for cancer prevention from vitiligo.

**Brief Summary:** Vitiligo skin harbors decreased somatic variation burden in cancer-associated genes and a concomitant augmentation in DNA repair response, explaining the lower incidence of cutaneous malignancies.

## Introduction

Multiple etiologies from auto-immune to genetic factors contribute to the pathogenesis of vitiligo, an acquired depigmenting disorder of skin that leads to loss of pigment producing melanocytes from the epidermis. While patchy depigmented appearance is the key and immediate worry for the subject, clinical management for photoprotection of the depigmented lesional skin is a serious concern^1^. This view emanates from observations on genetic depigmentary conditions such as albinism wherein complete loss of epidermal pigmentation poses a higher risk for cutaneous malignancies ^2,3^. Hence, it is believed that acquired depigmentation in vitiligo would pose a similar elevated risk. While in some studies the risk for cutaneous malignancies not to be significant in vitiligo subjects others report the enhancement in risk was found to be mildly elevated ^4–11^. Systematic evaluation of a large cohort of caucasian vitiligo subjects indicated a decreased risk for both melanoma and non-melanoma skin cancers. Analysis of independent retrospective comparative cohort in a multivariate 15 yearlong study reinforced this counter-intuitive observation ^6^. Thereby extrapolating from demographic studies, it is tempting to speculate that vitiligo could negatively influence either initiation or progression of cutaneous malignancies ^10^. Given the autoimmune etiology that targets melanocyte destruction, protection against melanoma could be rationalized, however a similar protection from non-melanoma skin cancer is perplexing. Therefore, these observations need to be substantiated with evidence at the tissue level and if/when validated, vitiligo would offer a tantalizing opportunity to understand molecular and cellular basis of cutaneous cancer prevention.

Skin being the largest organ of the body, is constantly exposed to environmental stress including the ultraviolet radiations that cause potentially deleterious mutations. Additionally, rapid clonal expansion of keratinocytes result in the progression of mutant cells into basal or squamous cell carcinomas (cBCC or cSCC)^12^. Accessibility of the tissue and recent advances in sequencing technologies make it feasible to map somatic clones in healthy skin tissue and provide valuable insights into the origin of mutations and map their progression to skin cancers. Deep-sequencing of selected panel of cancer associated genes suggests pervasive positive selection of somatic variations in normal, seemingly healthy skin ^13^. Notably, this seminal study reported that normal skin cells carry thousands of mutations, including oncogenic driver mutations observed in cancers and these are subject to strong positive selection. By performing a similar high-depth exome sequencing of pre-cancerous lesions and comparing it to adjacent sun exposed epidermis, the authors report that these cells may have already acquired 1-2 truncal driver variations ^14^. Further acquisition of additional alterations could promote parallel or linear progression to SCC. In another recent study the authors systematically studied somatic variations accrued in skin cell lineages and demonstrate that the amount and category of mutations in skin cells resemble many cancers ^15^. Sun-exposed skin cells have a higher mutation load attributable to UV exposure, but skin from hips protected by clothing have fewer variations that could be ascribed to UV ^15^. Thereby suggesting that the mechanisms that lead to carcinogenesis are functional in healthy skin cells and identification of not just the mutagenic triggers but also the cellular response to it would pave the path to decelerate the oncogenic progression.

In this current pilot study, we systematically map the somatic mutation burden using high-coverage whole exome sequencing in the matched lesion and non-lesional vitiligo skin derived from Indian subjects undergoing autologous transplantation. The somatic mutation trend observed was further investigated with respect to the altered vitiligo transcriptome, and tissue-level experimental validation in epidermal cells. Our analysis suggests that overall load and the clonal size of somatic variations in cancer-associated genes in the lesional depigmented vitiligo skin is significantly lower. Comparative skin transcriptome followed by validation from epidermal cells, highlights proliferation and DNA repair pathways to be augmented in the depigmented lesions. Thereby providing a possible mechanistic link to the lower prevalence of somatic variations in lesion despite the absence of protection by pigmentation. In tune with elevated DNA repair machinery, the lesional skin despite being exposed does not demonstrate UV-signature mutations. Thus, contrary to the expectations, and in sync with previous cancer incidence studies, our study establishes that the lesional exposed depigmented vitiligo skin does not have a high mutation burden and the extent of somatic cancer-associated variations is lower compared to the unexposed non-lesional pigmented skin. In future this study would pave the path to investigate mechanisms by which naturally altered states of tissue could provide meaningful strategies to prevent/reduce the risk of malignancies.

## Results

### Study design and analysis strategy

Clinical management of subjects with vitiligo lesions is two-pronged, targeting immune system to curb the autoimmune attack on melanocytes, and melanocyte repopulation to enable pigmentation of the patches. Targeting of immune system decelerates or altogether stops the progression of further depigmentation, resulting in disease stability. Depigmented patches that are recalcitrant to pigmentation but have remained stable for over a year are often managed by autologous epidermal-cell transplantation ^16–19^. After institutional ethics committee approval, informed consent was taken from these subjects for obtaining blood, lesional and surplus non-lesional skin after transplantation. Subjects chosen for this study were divided into three groups. Those willing to provide blood along with surplus non-lesional skin, and the lesional skin discard were included in the group I for deep-whole exome sequencing. We based our study on 18 exomes derived from a set of 6 subjects, from whom matched lesional vitiligo skin, non-lesional skin and blood were available for genome sequence comparisons. Individuals with only matched lesional and non-lesional skin were again divided into two groups II and III. The second set was taken for whole transcriptome analysis (4 subjects) and the third set of paired samples were reserved for tissue level, cell-based validation studies (3+3 subjects) (**Supplementary Table 1**).

Whole blood genome as reference allowed identification of baseline variations specific to the individual. This was followed by analysis of somatic variations in bulk, high-coverage whole exome sequencing from lesional and non-lesional skin. This strategy allowed pairwise comparisons for each of the six individuals. Ethical constraints precluded taking biopsies from adjacent peri-lesional area due to possible Koebner effect ^20^. Subjects with relatively exposed lesions uncovered by usual clothing style opted for transplantation thereby biasing the choice to sun-exposed lesional skin. Based on established protocols, non-lesional pigmented skin was chosen from the gluteal area which was relatively sun-unexposed (**Fig 1a** and **Supplementary Table 1**). From both locations split skin was taken during transplantation and prior to tissue processing dermal component was trimmed and the split skin containing predominantly the epidermis was taken for DNA or RNA isolation procedures. Enrolled subjects were of Indo-european ancestry, with a Fitspatric skin type IV/V residing in urban Indian city, either Delhi or Pune, exposed to environmental agents comparably.

**Figure 1:**
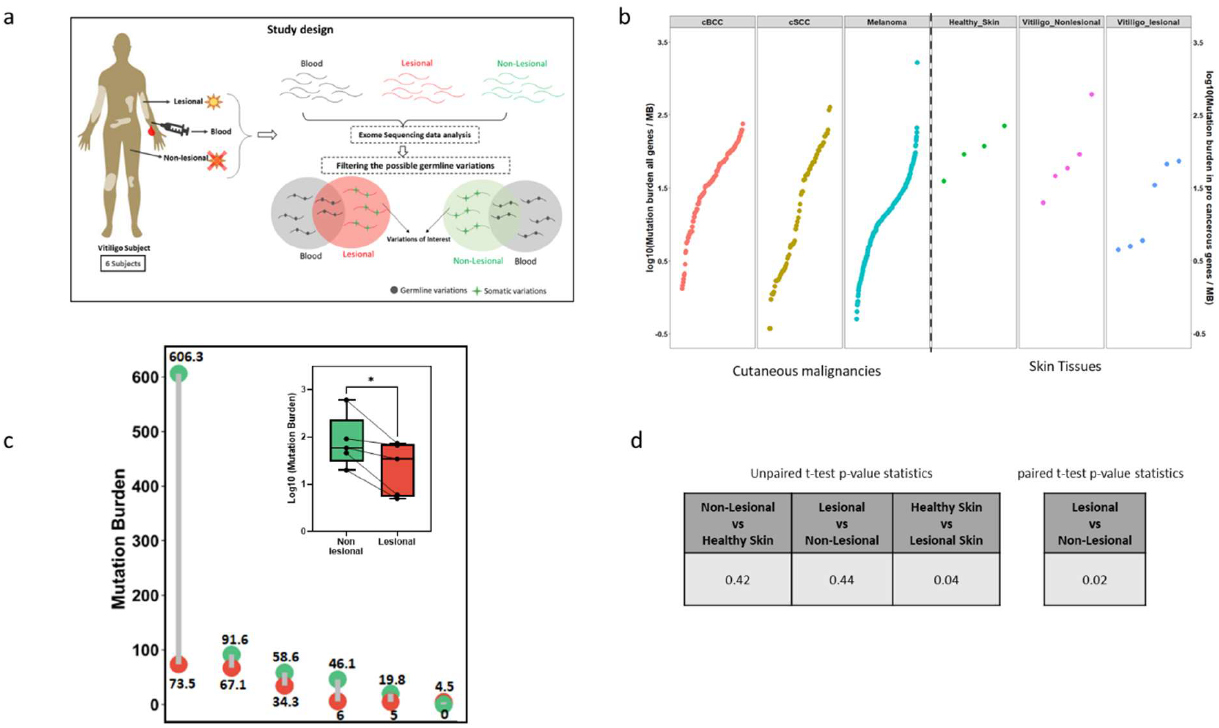
Spectrum of somatic variations across vitiligo skin tissues. (a) Study design for sample collection from vitiligo subjects and identification of somatic variations from group I of vitiligo subjects. (b) Rank order plot depicting the trend of variation burden across the three cutaneous malignancies, healthy skin and vitiligo lesional and non-lesional skin samples. The left Y-axis pertains to the three cutaneous malignancies and depicts Log_10_ mutation burden in all genes/Mb and the right Y-axis pertains to the three skin tissues and depicts Log_10_ mutation burden in cancer associated genes/Mb. (c) Dumbbell plot (with an inset Boxplot) depicting the mutation burden in lesional skin compared to non-lesional across 6 vitiligo patients. (d) Unpaired and paired t-test p-value statistics across healthy, non-lesional and lesional vitiligo skin comparison.

The somatic variation data generated from this study is qualitatively compared to that of the healthy skin from sun-exposed skin reported earlier ^13^. This study involved deep-sequencing of a selected exome of cancer-associated genes from multiple small skin punches and formed the basis of investigation for our current work. We resorted to bulk high coverage whole exome sequencing to account for pairwise differences between the lesional and non-lesional skin (**Supplementary Fig S1**). For the variation signature analysis, we resorted to the use of exome data from healthy skin from Moore et. al. for which the whole exome sequencing data was available ^21^. Sun exposure, location of the tissue as well as differences in ethnicity, remain potential confounders for interpretation. Comparison with relevant malignancies namely cutaneous basal cell carcinoma (cBCC) and cutaneous squamous cell carcinoma (cSCC) which are cancers arising out of keratinocyte lineage and malignant melanoma arising from melanocytes were included from cBioportal ^22,23^. While these studies were compared by metanalysis, the key comparison is between the matched vitiligo lesional and non-lesional exome in cancer associated genes.

### Burden of somatic variations in vitiligo skin

Possible sequencing artefacts were removed by applying stringent quality and depth filters. The germline variants were then filtered by comparing to their matched blood exome (Details in methods section). A total of 76,310 and 20,202 somatic variations were detected in the non-lesional and lesional vitiligo skin tissue respectively. These were unique to skin and not detected in the corresponding blood samples of vitiligo subjects **(Supplementary Fig S2-S6)**. Interestingly, the variability across the non-lesional tissues within individuals was higher compared to lesions, which had lower but uniform number of variations.

At the tissue level, mutation burden in oncogenes would prove prognostic to objectively assess the “cancer-risk” and explain observations on demography-based incidence of non-melanoma skin cancers among vitiligo subjects. Based on the currently emerging trend from various studies, healthy skin harbors mutations in cancer-associated genes, and these are under positive selection. The burden of mutation in healthy skin was found to be lower, but in overlapping range of cutaneous malignancies ^13^. We estimated the range of mutation burden in both vitiligo skin tissues and compared with the healthy skin as well as cutaneous malignancies **(Fig 1b)**. While estimates for cutaneous malignancies were based on whole genome mutation data from cBioportal, comparable analysis method could be employed for healthy skin as reported earlier ^13^. We observed that the mutation burden in the cancer-associated genes in vitiligo lesion is lower compared to the healthy skin as well as the non-lesional vitiligo skin **(Supplementary Table 3-5)**. Missense variations were absent in majority of the lesional vitiligo tissues and non-lesional samples had barely 1 to 3 variations **(Supplementary Table 6 & Supplementary Fig S7)**. Lesional skin being depigmented and sun exposed, has UV as an external source that could cause somatic variations, whereas the matched non-lesional skin is pigmented and sun protected. Despite this the anticipated trend of more variations was not observed in vitiligo lesions. Of the many possibilities, this could be indicative of certain endogenous factors responsible for providing protection which are either absent/inefficient in healthy and non-lesional pigmented skin tissues. The mutation burden in the healthy skin and non-lesional tissues were comparable albeit differences in sampling and this needs to be further explored by systematic comparative studies with larger sample size. The pair-wise comparison of vitiligo skin samples reveals that lesional skin indeed have a lower mutation burden in 5 out of 6 subjects despite being diseased and sun-exposed (**Fig 1c**). One individual VD31, did not follow the trend had 0 somatic variations mapping to cancerous genes in the non-lesional skin. The difference was observed to be statistically significant based on paired t test analysis **(Fig 1d)**.

We investigated whether the increased number of variations in non-lesional tissue arises from majority of cells carrying the same alternate allele (clonal), or a small subset of cells carrying the alternate allele (subclonal) within the sampled tissue. This was carried out using the variation data as described previously (with modifications, in the methods section). Towards this we estimated variant allele fraction (VAF), which is a measure of clonal/subclonal population of cells within a tissue ^24^. VAF of 1 is indicative of clonal cell population with all cells harboring the alternate allele. Owing to differences in sequencing technologies and single data collection per sample, our analysis of VAF would capture only bigger clonal representation in the skin. We estimated that the minimum detectable clone size, in the 1cm^2^ of tissue with an average depth of 500x and the minimum of 20 reads supporting the alternate allele would be 8mm^2^ or more. As the aim of the analysis is comparison of lesional to non-leisonal skin, this was deemed sufficient. Furthermore, in our analysis germline variations are removed, hence the estimates of clone-sizes are not directly comparable with the already published healthy skin data. The overall pattern of distribution of VAF across lesional and non-lesional tissues were comparable **(Supplementary Fig S8)**. Presence of higher proportion of somatic variations in non-lesional skin is indicative of bigger clones compared to the lesional tissue.

### Size distribution and clonal analysis of cancer associated gene variations is lower in vitiligo lesions

We then proceeded to investigate whether higher mutation burden in healthy and non-lesional skin correlates with clonal size in the cancer driver genes. Towards this we created a circlepack plot. Herein the outer blue, red and green circle represents the tissue type, the inner white circle denote genes and grey circle represents variations, size of which is proportional to the clone size estimated by VAF. It is important to note here, since the data pertaining to the vitiligo samples were generated in the same experiment, size of the grey circle in the non-lesional and lesional tissues are comparable **(Fig 2 a & b)**. We observe bigger clones in non-lesional tissue compared to the lesional skin, that had fewer smaller clones and the clonal representation per gene was also observed to be low. We argued that one of the non-lesional skin had almost 10-fold higher somatic mutation burden in cancer genes and this could skew the size distribution. A reanalysis of the data with omission of VD32 resulted in similar observation. Therefore, we conclude that the number of variations and their overall VAF in the lesional skin were comparatively lower than the non-lesional skin **(Supplementary Table 7)**.

**Figure 2:**
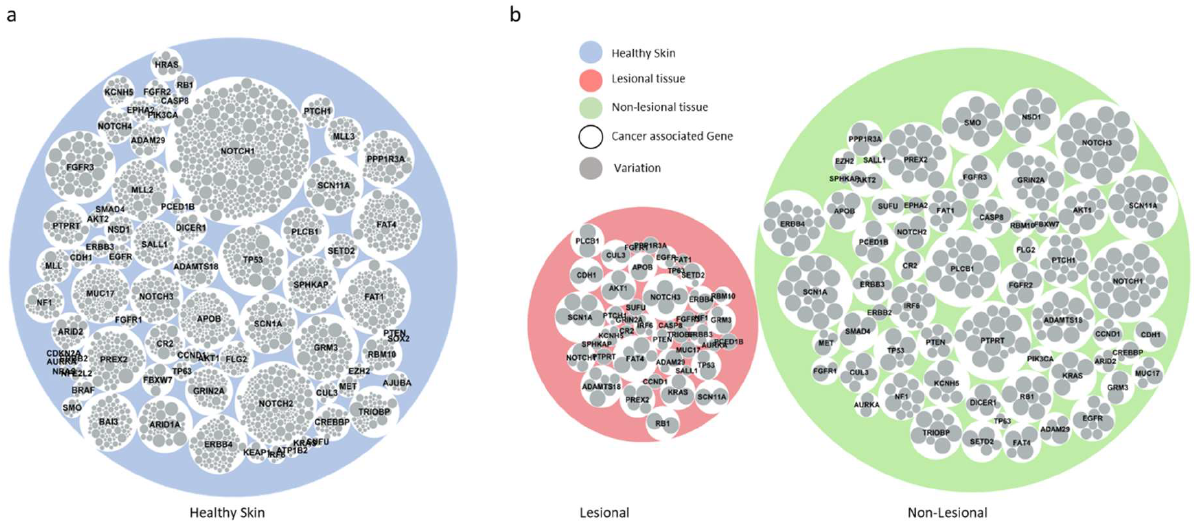
Lower clonal variations in matched vitiligo skin. Circlepack plot depicting the clonal/subclonal variations across the lesional and non-lesional vitiligo skin (a) Healthy skin data published elsewhere (b) non-lesional and lesional vitiligo skin. The outer coloured circle (blue, red, green) represents the tissue type; The white circles depict the cancer associated genes; Innermost grey circle depicts variations. The size of the grey circle is proportional to the VAF value in their respective tissue type.

### Transcriptional footprint in lesional vitiligo skin suggests increased cell proliferation

The smaller clones, lesser mutation burden in cancer genes in lesional skin are strongly interrelated and explain the lower incidence of non-melanoma skin cancers in vitiligo subjects. However, insights into the possible mechanisms that result in this reduced mutation burden would add immense value to our understanding of the initiation and progression of cutaneous malignancies. Towards this end, we performed whole transcriptome sequencing of paired lesion and non-lesional skin tissue from 4 vitiligo subjects from Group II (**Supplementary Figure S9**). The observations from vitiligo transcriptome were comparable to our previous microarray results from 15 vitiligo subjects ^25^. We observed that pathways pertaining to melanogenesis is downregulated in the lesional tissue, which could be attributed to the loss of mature melanocytes **(Supplementary Table 8)**. Pathways upregulated in the lesional tissue could either be the underlying cause of depigmentation or could possibly explain the response of skin to the absence of pigmentation. We observed an upregulation of nuclear division and cell cycle related processes among the top upregulated pathway terms **(Fig 3a and Supplementary Table 8)**. To validate this observation, we performed immunohistochemistry for two key proliferation-associated markers *MCM6* and *PCNA* (skin tissue from Group III subjects). Across the three individuals, immunohistochemistry analysis showed higher staining of these two markers. Flow cytometry-based cell cycle analysis of epidermal cell suspension corroborated these findings and we observed higher number of cells in S+G2+M stages of the cell cycle, suggesting more number of proliferative cells in the lesional epidermis. **(Fig 3 b&c)**. Furthermore, several of the genes pertaining to cell proliferation *(CCNE2, CDK2, CCNA2, CDC2, KI67, CDC7)* normalized to 18srRNA were upregulated upon RT-PCR validation **(Fig 3d)**. In our earlier work, we had systematically investigated architectural changes in the epidermis that support our observation on higher proliferation of epidermal cells ^25^. Thereby, we unequivocally establish increased proliferation in lesional vitiligo skin. Higher proliferation could facilitate faster turnover of mutant clones and possibly explain the observed lower burden. Whether the possible fast epidermal turnover is sufficient to keep a check on the mutation burden remains speculative.

**Figure 3:**
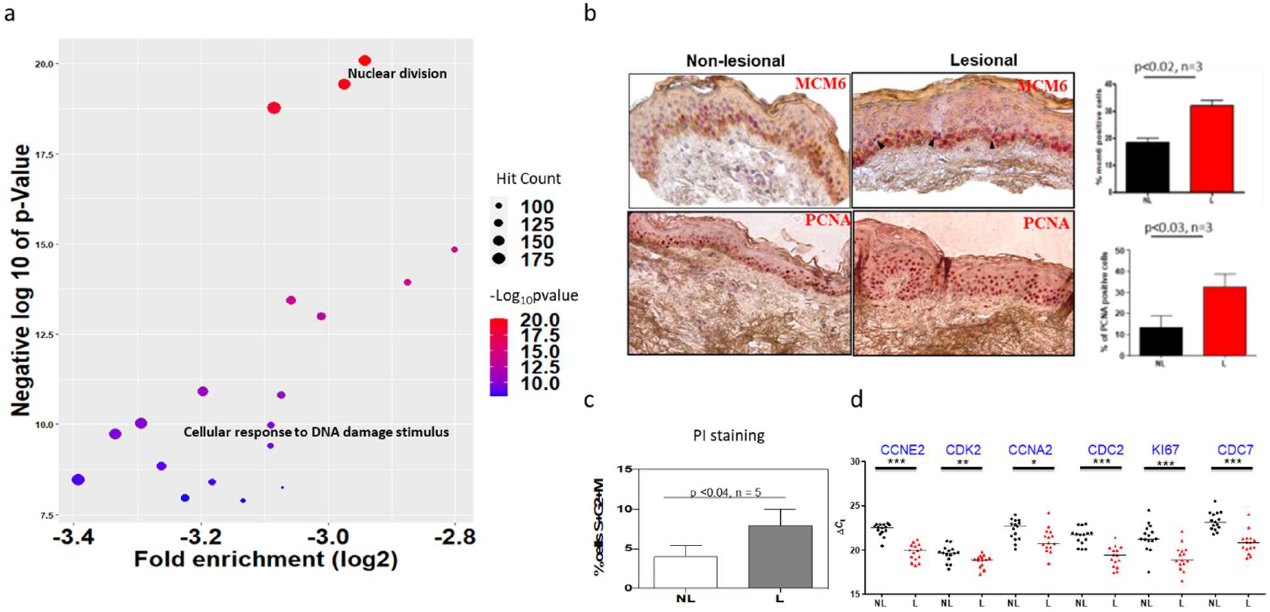
Analysis of transcriptomic footprint in the two matched vitiligo skin. Transcriptomic analysis from matched lesional and non-lesional vitiligo skin from group II subjects. (a) Bubble plot depicting enriched processes in the upregulated set of genes, based on whole transcriptome sequence analysis of lesional wrt non-lesional vitiligo skin. (b) Representative images and quantitation of immunohistochemical staining for MCM6 and PCNA. (c) Cell cycle analysis based on propidium iodide staining of epidermal cells. Bar plot depicts number of cells in S+G2+M phase of cell cycle. Paired students t test results are depicted. (d) RT-PCR results showing upregulation of proliferation related genes.

### Elevated DNA damage response explains the lower mutation burden of lesional vitiligo skin

On a closer analysis of the upregulated pathways, DNA repair response emerged as a probable link that could explain the observed somatic variation burden of the lesional skin. Based on two-way hierarchical clustering of expression values in the heat map, expression of a set of DNA repair genes is sufficient to segregate lesions from the non-lesional samples. Suggesting that significant differences exist in the expression of DNA repair genes. Most of these genes were found to be upregulated in vitiligo lesions **(Fig 4a, Supplementary Fig S10a)**. To functionally validate alterations in DNA repair capacity we performed comet analysis from epidermal cell suspension that would be predominantly populated by the epidermal keratinocytes. The single cell electrophoresis followed by propidium iodide (PI) staining resulted in comets, and quantification of tail moment suggests that lesional skin has lower amount of DNA breaks than the non-lesional skin **(Fig 4b)**. Lower number of breaks could be due to better resolution of DNA damage signifying ongoing repair. Since alterations in cell cycle was also observed, a concomitant increase in DNA repair pathways strongly suggests a replication coupled repair response. Substantiating this possibility, in a volcano plot of replication coupled-DNA repair pathway, most of the genes were significantly upregulated in the lesional vitiligo skin **(Fig 4c, Supplementary Fig S10b)**.

**Figure 4:**
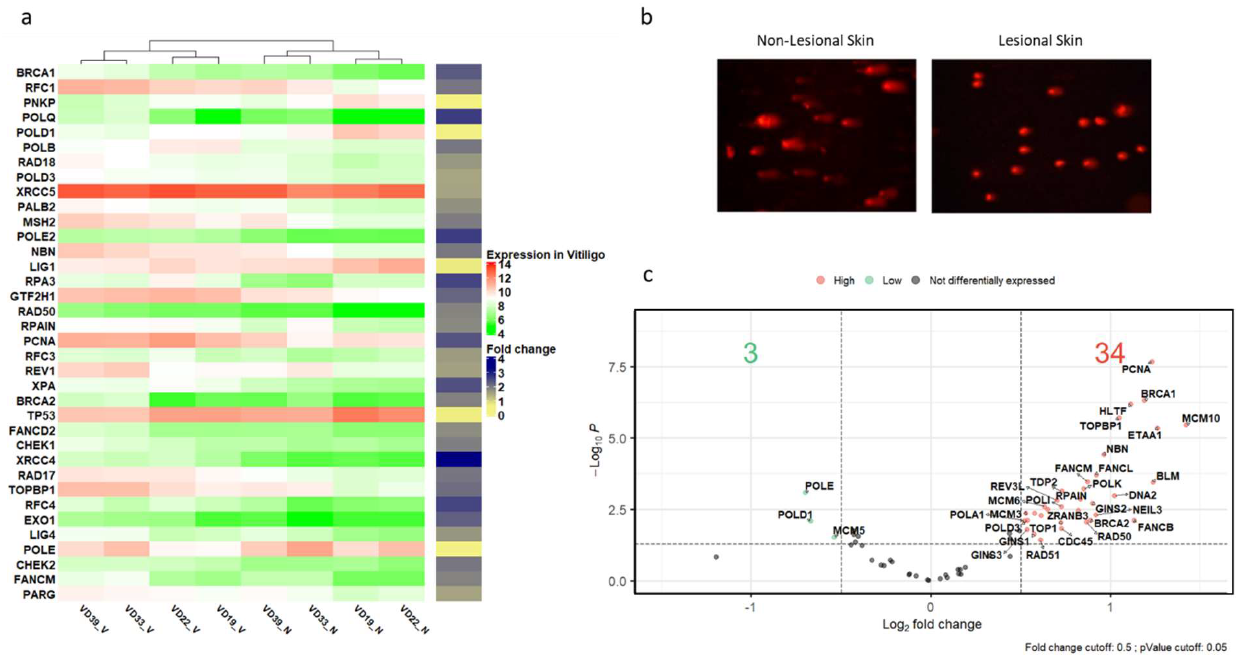
Increased DNA damage and repair in lesional skin tissues is through replication coupled DNA repair. (a) Heatmap depicting the expression pattern of DNA damage genes in group III vitiligo samples. The green-red color represents the normalized expression values and the yellow-blue annotation bar is the fold change in lesional wrt non-lesional vitiligo skin for that gene. (b) Comet assay depicts DNA breaks in lesional skin tissue compared to non-lesional skin. (c) Volcano plot for genes pertaining to replication coupled repair pathway. Here red dot represents a gene to be significantly up in Lesional sample whereas, green dot represents significant downregulation, based a p value cut off of 0.05 and a Log2 fold change cut off of 0.5.

A recent study provides a foot-print of the three clonogenic pools of epidermal keratinocytes that are responsible for maintenance of the human epidermis. Holoclone-forming cell has hallmarks of stem cells and are bestowed with higher self-renewal capacity as well as their ability to repair DNA damage ^26^. Interestingly, we observed that the signature of holoclones closely parallels the observations made in vitiligo. The set of holocone signature was majorly found to be upregulated in vitiligo lesions and segregated the lesions and non-lesional transcriptome in both RNA-sequencing reported in this study as well as the microarray-based results from a larger number of individuals reported earlier (**Supplementary Fig S11**). Thereby, it is likely that the lack of melanin in lesional skin and associated changes elicit a response via replication coupled repair in keratinocytes which could account for the lower mutation in vitiligo.

### Mutational signature analysis and comparison across cutaneous malignancies reveal a distinct pattern

Lower mutation burden and smaller clone size in the lesional vitiligo tissue suggests an underlying difference in somatic mutagenesis. To identify the footprint of the somatic mutations, we performed mathematical extraction of existing COSMIC signatures in the skin tissue data from healthy, vitiligo skin tissues and the three cutaneous malignancies derived from published work ^27–34^. As reported earlier, UV related signatures (SBS7a&b) were the top signatures in skin cancers as well as the healthy skin with a median cosine similarity of 0.94 ^21,35^. However, to our surprise UV signature were not present not only in the unexposed non-lesional skin, but also in the sun-exposed lesional tissue. Instead, SBS5 signature (cosine similarity > 0.86) were the only signatures which was significantly enriched in both lesional and non-lesional vitiligo tissues. SBS5 was also found in cBCC and healthy skin however, their cosine similarity was not significant (0.72 and 0.54) (**Fig 5 a, Supplementary Table 9**). The number of variations contributing towards SBS5 signature were higher in non-lesional tissues compared to the lesional tissue, in 5 out of 6 patients (**Fig 5b, Supplementary Fig S12**). SBS5 characterized by C>T and T>C transitions, has been shown to positively correlate with age and tobacco smoking. However, its etiology remains unexplored and debatable ^35,36^. Both C>T and T>C transitions are observed in vitiligo skin prominently (**Fig 5 c**). We speculate that T>C could be attributed to dark CPD formation as an earlier prominent work identified TG lesions in melanocytes ^37^. In the light of altered cell-proliferation coupled DNA repair pathway activation, it is likely that despite being exposed and depigmented, the lesional vitiligo skin harbors lower burden of SBS5 signature mutations and this could also explain the absence of UV-signature mutations. However, a systematic comparison with healthy individuals of similar ethnicity and skin type may be needed to make this comparison feasible.

**Figure 5:**
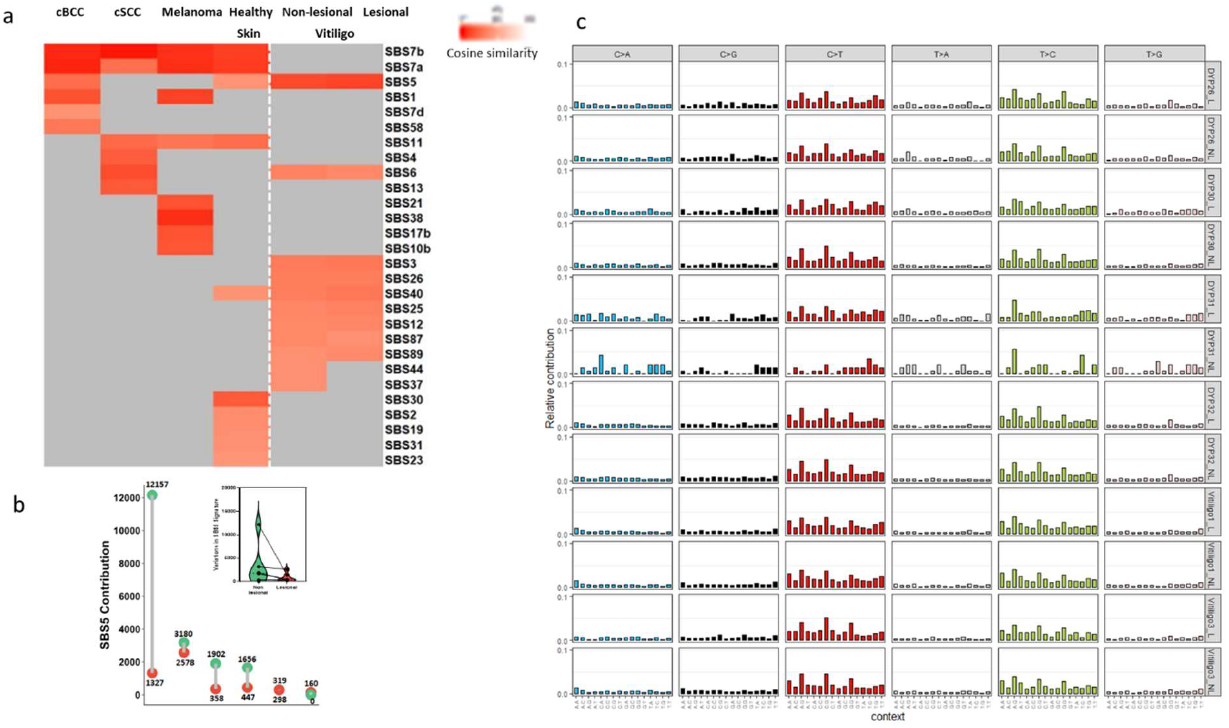
Mutation signatures in lesional and non-lesional skin. (a) Heatmap of the cosine values for the extracted signatures across multiple skin cancers, healthy and vitiligo skin samples. (b) Dumbbell plot (with an inset Boxplot) depicting the number of variations contributing to the SBS5 signature by lesional and non-lesional skin. (c) The relative contribution of 96 mutational profile across vitiligo skin samples.

## Discussion

The current genomic exploration in vitiligo reveals lower somatic variation burden that endorses the earlier demographic observations on decreased incidence of skin cancer among vitiligo subjects. It is intriguing that rewiring of gene expression triggered by the disease, renders a state that offers protection from a more detrimental skin condition. In this regard the lesional vitiligo skin appears to be an enantiostatic state wherein the skin cells rewire and adopt an alternate steady state ^25^. Since in this pilot study the vitiligo skin is sun exposed, it is likely that the observed differences could be mediated by the influence of environmental UV exposure on the depigmented skin. In future, detailed large-scale comparison that account for age, skin type, gender, sun-exposure as well as matched samples across subjects including healthy volunteers free of the vitiligo is warranted. Thereby, our initial exploration does indicate towards an altered state, and raises further questions that would need to be systematically investigated in greater detail with more elaborate comparisons.

Human epidermis contains long-lived quiescent stem cells, as well as cells with varying proliferative capacities. Though these have been known for a very long time, their molecular identity has only emerged recently. The clonogenic keratinocytes are contributed by holoclones, meroclones, and paraclones that appear to harbour distinct functional role in skin ^26^. While meroclones represent the bulk of epidermal clonogenic population, holoclones appear to assume the stem cell function and are likely to come to play during epidermal regeneration. However, in vitiligo lesions DNA damage could be efficiently repaired in the expanded holoclone population but could be present as somatic variations in the minor population of slow-cycling meroclone or paraclones that do not have augmented DNA repair machinery. These possibilities and the detailed underlying mechanisms need to be investigated in the future.

Despite, vitiligo lesions accounting for only around 20-30% of body surface area on an average the observed decrease in cancer incidence is seemingly more pronounced from the demography-based incidence data. Thereby, it is tempting to speculate that the non-lesional skin may be different from the normal skin of a healthy individual and could contribute towards cancer protection. This possibility is further endorsed by the absence of UV-signature variations in the non-lesional skin, while the same is reported from the sun-unexposed healthy skin^15^. However, this is to be interpreted cautiously, as the healthy skin is from the caucasian subjects which is significantly different from the fitzpatrick type IV/V skin analyzed in our study. The lesional vitiligo skin does not harbour UV-signature mutations, implicating the role for vitiligo specific alterations to override ethnic difference of skin types. This would need to be formally explored in a large-scale comparison involving healthy individuals along with vitiligo subjects from the same ethnic background.

While UV signature could not be detected, SBS5 emerged to be significant. In this signature ubiquitous metabolite mutagen-induced modification of DNA is proposed to accumulate. Earlier studies observed this signature in kidney cells that are constantly exposed to these metabolites ^36^. Though SBS5 positively correlates with age, we observed this signature across individuals of all age groups. In the non-lesional pigmented vitiligo skin, such signatures could arise due to UV-induced modification of DNA^37^. Whereas in the lesional skin due to efficient repair mutations contributing to SBS5 are substantially reduced.

Evolution of pigmentation is adaptive, and this feature is evident in the naked human skin wherein pigmentation provides protection from UV-damage ^38^. The loss of pigmentation in acquired disorders is strongly believed to render a maladaptive state to the skin. This belief is reinforced by depigmenting genetic disorders such as albinism that result in skin cancers. Vitiligo seems to provide protection from UV-induced mutagenesis by an alternative mode, involving replication coupled DNA repair pathways. Hence, pigmentation though widely employed by nature, is not the sole mechanism for photoprotection and this could be alternately tackled by modulating the cell-state. Thereby vitiligo provides a tantalizing opportunity to understand this state of skin in greater detail. While vitiligo could offer protection from somatic mutations, photoprotection may still be required to prevent sun burn ^10^. Our initial insight therefore needs to be investigated further to derive precise mechanisms and identify key actionable players that could be translated not only for cancer prevention, but also to promote desirable mutations in skin cells to revert genodermatoses.

## Acknowledgement

Council for Scientific and Industrial Research (CSIR), India, through grants TOUCH-BSC0302 and RegenX-MLP2008 provided support to the execution of this work.

## Materials and Methods

### Ethics statement and Sample collection

All the patients enrolled in this study were of North Indian ancestry and were not undergoing any systemic/topical treatments at the time of sample collection. Split skin from lesional and non-lesional epidermal tissues were collected from six Vitiligo subjects (Group I). Matched whole blood was also collected to filter the germline variants. For whole transcriptome sequencing and validation studies matched lesional and non-lesional samples were collected from 4 and 3 vitiligo subjects (Group II and III). The stable lesional patches for at-least 6 months were collected from the sun-exposed affected areas and the non-lesional tissues were collected from the sun protected gluteal region. Institutional Human ethics committees of National institute of Immunology, New Delhi and CSIR-IGIB approved the study (IGIB/IEHC/11; Dated:30.08.2012), and it is in agreement with Declaration of Helsinki principles. All the specimens collected in this study were undergoing skin grafting procedure and were taken after informed consent from the patients **(Supplementary Table 1)**.

### Handling of skin samples

Skin samples were collected under aseptic conditions in transportation medium comprising Hanks’ Balanced Salt Solution (HBSS) and 10X antibiotic-antimycotic. The samples were preserved in cold till the processing. The skin was cut into smaller pieces of approximately 0.5 cm and additional connective tissues were removed. The tissue samples were sterilized by alternate washes in 70 % ethanol and transportation medium.

### Reporting of data

Statistical methods were not deployed to pre-determine the size of the samples for this study. The experiments were not randomized, and the investigators were not blinded to allocation, either during the experimentation or during the assessment of outcome.

### Whole exome sequencing and data analysis

The DNA isolation protocol was adapted from Rani, R. et al ^39^. The whole exome sequencing of the samples was carried out using Nextera Rapid Capture Expanded Exome kit. The libraries were prepared according to the manufacture recommended protocol and 100 bp paired end reads were generated on HiSeq2000 (Illumina Inc, USA). Paired-end sequencing of the isolated DNA from all the matched lesional, non-lesional and blood tissues was performed at CSIR-IGIB New Delhi. To increase the depth of coverage each sample was run on 6 lanes and were merged in the later steps of the pipeline to generate single file per tissue per sample. Initial quality control and filtering out the bad quality reads were done using FASTQC and Trimmomatic ^40,41^. The high-quality sequenced reads were then aligned to human reference genome (NCBI build GRCh38 ^42(p38)^) using BWA-Mem ^43^. Post alignment the PCR duplicates were removed, and the multiple lane files were merged using picard tool. The variants were called using GATK’s Haplotype caller (version 3.7) and annotated using Annovar ^44^ resulting in the VCF file for all tissues from all patients **(Supplementary Fig S2)**.

### Post VCF quality control and removal of germline variations

A depth cutoff of >=20, genotype quality >= 30 and quality >=20 was applied to filter the VCF files and remove any possible sequencing artifacts ^45^. Since, the main objective of this study was to identify the patterns/processes (not inherent in nature) that provides protective mechanism against non-melanoma skin cancers in Vitiligo subjects filtering out any possible germline variations is one of the crucial steps. To accomplish this, we leveraged the matched blood tissue and compared it to lesional and non-lesional tissue. Any variant which came common with blood in lesional/non-lesional tissue was considered as germline variant and was removed from the downstream analysis **(Supplementary Fig S2 & S3)**.

### Variant allele fraction estimation

We calculated Variant allele fraction for the variations. Variant allele fraction (VAF) is an estimate of clonal and subclonal variations, it is calculated as the ratio of reads supporting the alternate allele to that of total number of reads at that particular position (depth) ^24^.

### Integrative analysis with Healthy skin and Melanoma, Non-melanoma skin cancers

Healthy skin and melanoma and non-melanoma skin cancer datasets:

We leveraged the high-depth sequencing data for healthy skin samples from Martincorena, Iñigo, et al. (2015) ^13^. This dataset comprises of sun-exposed eyelid skin samples from 4 individuals (234 biopsies) undergoing blepharoplasty. The coding exons of 74 genes implicated in cancers were sequenced with an average coverage of 500x.

For the three cutaneous cancer types, data was downloaded from cbioportal (Dated:**4.08.2021**) ^22,23^. Melanoma dataset comprising of 346 subjects were included from Akbani, Rehan, et al. (2015) ^30^. The Cutaneous Basal cell carcinoma from 126 samples was taken from Bonilla, Ximena, et al. (2016) ^31^. Cutaneous Squamous cell carcinoma dataset comprises of 151 subjects from multiple studies ^32–34^.

### Estimation of mutation burden across datasets

We adapted the similar method to estimate mutation burden as described by Martincorena, Iñigo, et al. where ^13^:

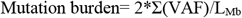

VAF is the variant allele fraction and L_Mb_ is the length of the bases sequenced in megabase (MB). L_Mb_ for Martincorena, Iñigo, et al. data and Vitiligo dataset is 0.67MB. Beacuse, Martincorena, Iñigo, et al. performed targeted sequencing on 74 cancer causing genes we therefore, filtered our dataset with respect to these genes only to minimize under-estimation of mutation burden and maintain consistency across the analysis ^13^.

For the cancer datasets, owing to the unavailability of allele fraction information and for the consistency across the analysis we estimated mutation burden as n_exome_/30MB, as described in Martincorena, Iñigo, et al. ^13^ (**Supplementary Table 6**)

### RNA sequencing and data analysis

The RNA isolation was performed using the protocol mentioned by Singh, Archana, et al. ^25^. The RNA quality check was performed using Agilent 2100 Bioanalyzer system before library preparation. The RNA libraries were prepared using Illumina’s TruSeq total RNA Sample preparation Kit according to the manufacture’s guidelines. The sequencing was performed on Illumina HiSeq2000 using Truseq PE cluster kit and 100bp paired end reads were generated. Paired-end sequencing of the isolated RNA from all the matched lesional and non-lesional tissues was performed at CSIR-IGIB New Delhi. RNA-seq data was initially checked for the quality and the bad quality reads were filtered using FASTQC and Trimmomatic ^40,41^. Post quality control and data trimming, high quality reads were aligned against the human reference genome (NCBI build GRCh38 ^42(p38)^) using STAR aligner ^46^. Post alignment reads were quantified using HTSeq ^47^ and then the data was checked and corrected for any batch effects using SVA and PVCA R packages ^48,49^. The data was normalized and differential gene expression was calculated using DESeq2 R package ^50^. ToppGene Suite was used for estimating the biological processes that were enriched in the Vitiligo dataset ^51^.

### Immunohistochemistry for MCM6 and PCNA

Skin biopsies were fixed with 10% formalin, dehydrated, and embedded in paraffin. Sections (4 µm thick) were placed on polylysine-coated slides, deparaffinized, hydrated, antigen retrieved with 10mM citrate buffer pH 6.0, and blocked with protein block serum free (Dako cytomation). Slides were stained with rabbit polyclonal antibodies to PCNA (dilution 1:700; abcam) or MCM6 (1:100 dilution, abcam) for 30 minutes at room temperature followed by incubation with labelled polymer alkaline phosphatase (Dako Cytomation) for 30 minutes at room temperature. Reaction was developed with fast red and counterstained with hematoxylin.

### Cell Cycle Analysis

Epidermal cell suspension from non-lesional and lesional skin was fixed with 1ml ice cold 70% ethanol added dropwise, and the cells were incubated at 4 °C overnight. The cells were then resuspended in 500μl staining solution containing 20 μg/ml Propidium Iodide, 15 μg/ml RNAse A and 0.05% triton X 100, incubated for 60 min at 4°C, and analyzed by flow cytometry (BD LSR).

### qRT-PCR and analysis

Total RNA was isolated from non-lesional and lesional epidermis using the Qiazol reagent (Qiagen) and converted into cDNA (1^st^ strand cDNA synthesis kit, Takara). Real time PCR was carried out using the LC480 Thermocycler (Roche) using SYBR Green chemistry (Fermentas Intl, Ontario, Canada) as per the manufacturer’s instructions. The Ct method (Δ Ct value) was used for gene expression analysis. Ct values were normalized to their corresponding 18S rRNA levels to obtain Δ Ct values. The reactions were performed in triplicates, and Δ Ct values differing by an SD of Δ 0.5 between replicates were eliminated from further analysis. Fold change was calculated using 2^(-Δ Δ Ct), where Δ Δ Ct represents Δ Ct of lesional–Δ Ct non-lesional epidermis. Paired Student’s t-test was performed to study the significance of the Δ Ct value pairs. All statistical analyses were carried out using the Graph-Pad Prism software (version 5, San Diego, USA) and p values <0.05 were considered statistically significant.

### Alkaline comet assay

50 μL suspension of 30,000 cells was added to 550 μl of 0.75 % of low melting agarose. Single cell suspensions of these cells were loaded on 24 × 50 mm glass slides pre-coated with 0.1 % low melting agarose. The cell suspensions are allowed to cool for 2-3 min on ice. The slides were then dipped into Alkaline lysis buffer (NaCl-2.5 mM, EDTA 100 μM, Tris base 10 mM, Sodium sarcosinate 1 %, DMSO 5%) for 20 min at 4 °C. The slides were then washed in MilliQ and again incubated in alkaline electrophoresis buffer (NaOH 0.3 M, EDTA 3 mM) at 4 °C for 40 min. The slides were then run in freshly prepared alkaline electrophoresis buffer at 2 V/cm and 300 mA for 20 min. Finally, the excess alkali was neutralized in a neutralization buffer (Tris base 0.4 M) for 5 min. The slides were then dried at 50 °C and kept at 4 °C overnight. The slides were then rehydrated in milliQ for 5-7 min and stained with 50 μM of propidium iodide stain.

Images were obtained in a Leica fluorescence microscope at 40X magnification. Mean tail length was measured for 50 random cells (25 from each replicate) and analyzed using Komet 6.0 (ANDOR technology, Belfast, UK) software, as per guidelines.

### Estimation of mutational signature

For the estimation of the optimal contribution of previously defined COSMIC mutational signatures in our dataset we used signature refitting function of MutationalPatterns R package ^27–29,52^. For the cancer dataset, the signatures were estimated using MutationalPatterns and Maftools R package ^52,53^. A cutoff of > 0.85 cosine similarity value was considered as significant ^54^. All the sample meta information is provided in supplementary table 3-5. The healthy skin dataset from 2 subjects was taken from Moore, Luiza, et al. (2021) and the mutational signatures was estimated by the same method described above_21_.

### Statistical analysis

GraphPad Prism (8.0.1), R (Version 4.0.3) and Microsoft excel was used for statistical analyses and plotting of the graphs. P-value less than 0.05 were considered statistically significant throughout the analysis.

---

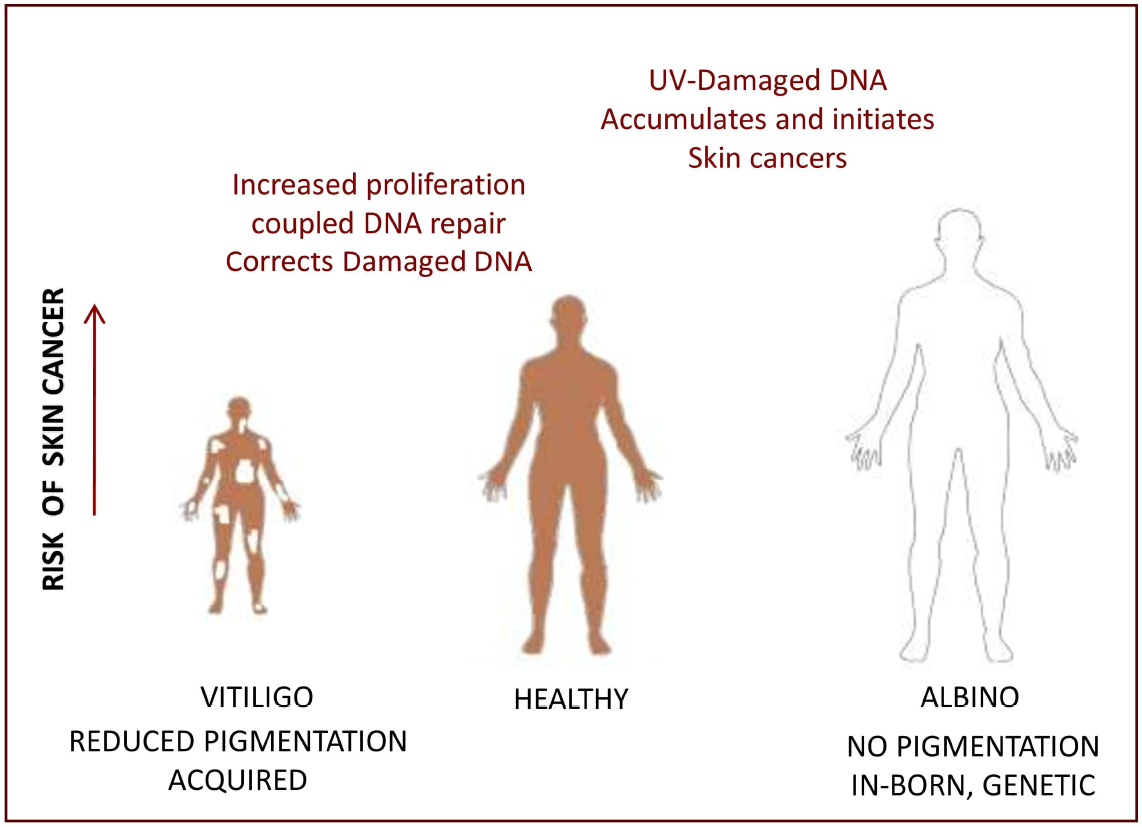

