## Supplementary Figures for "Whole exome-sequencing of vitiligo lesions indicate lower burden of somatic variations: implications in risk for non-melanoma skin cancers"

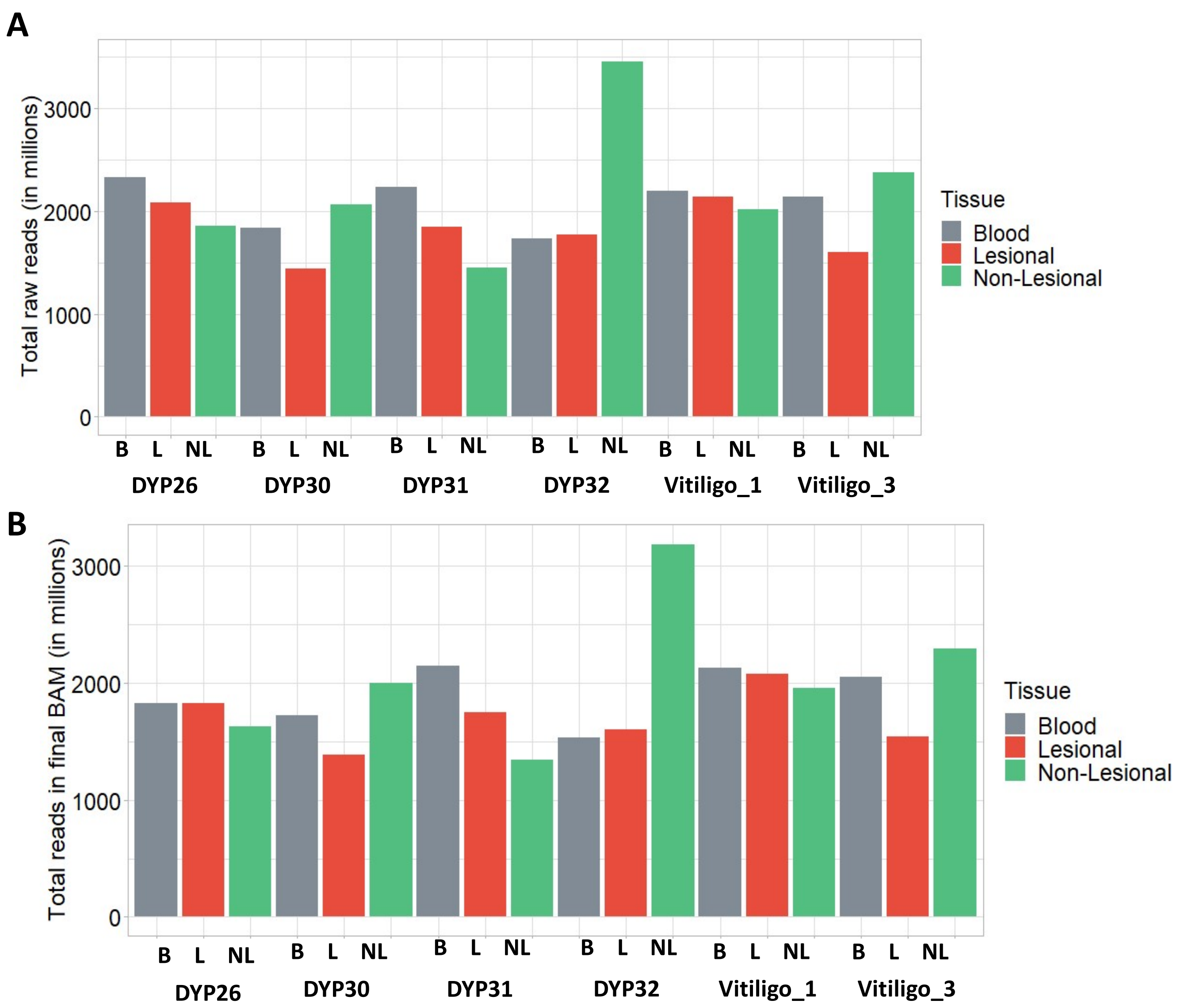


**Figure S1: Distribution of number of reads across samples (A) In raw fastq file & (B) In final merged BAM files**

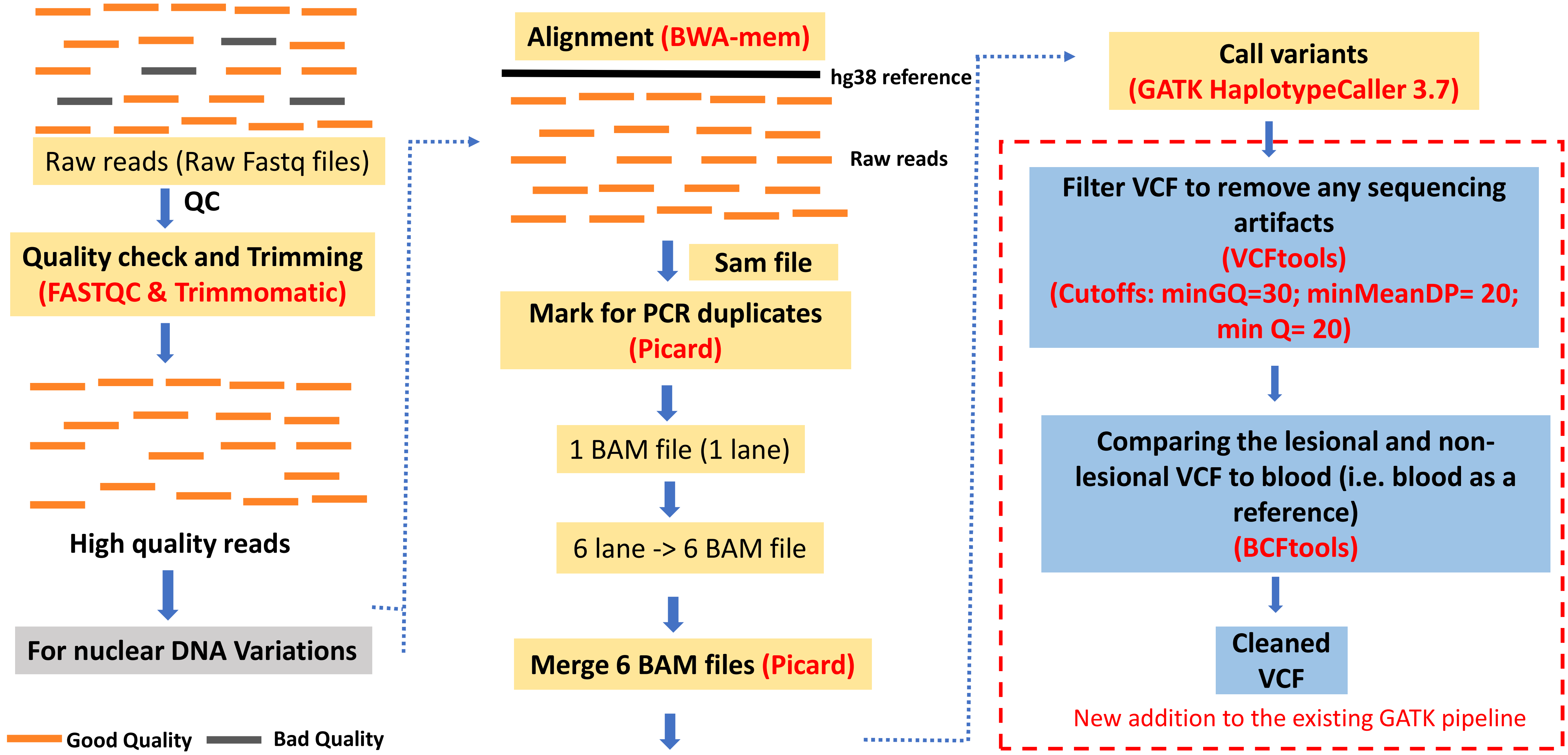


**Figure S2: Workflow followed for the whole exome sequencing data analysis: From Fastq file till final VCF generation**

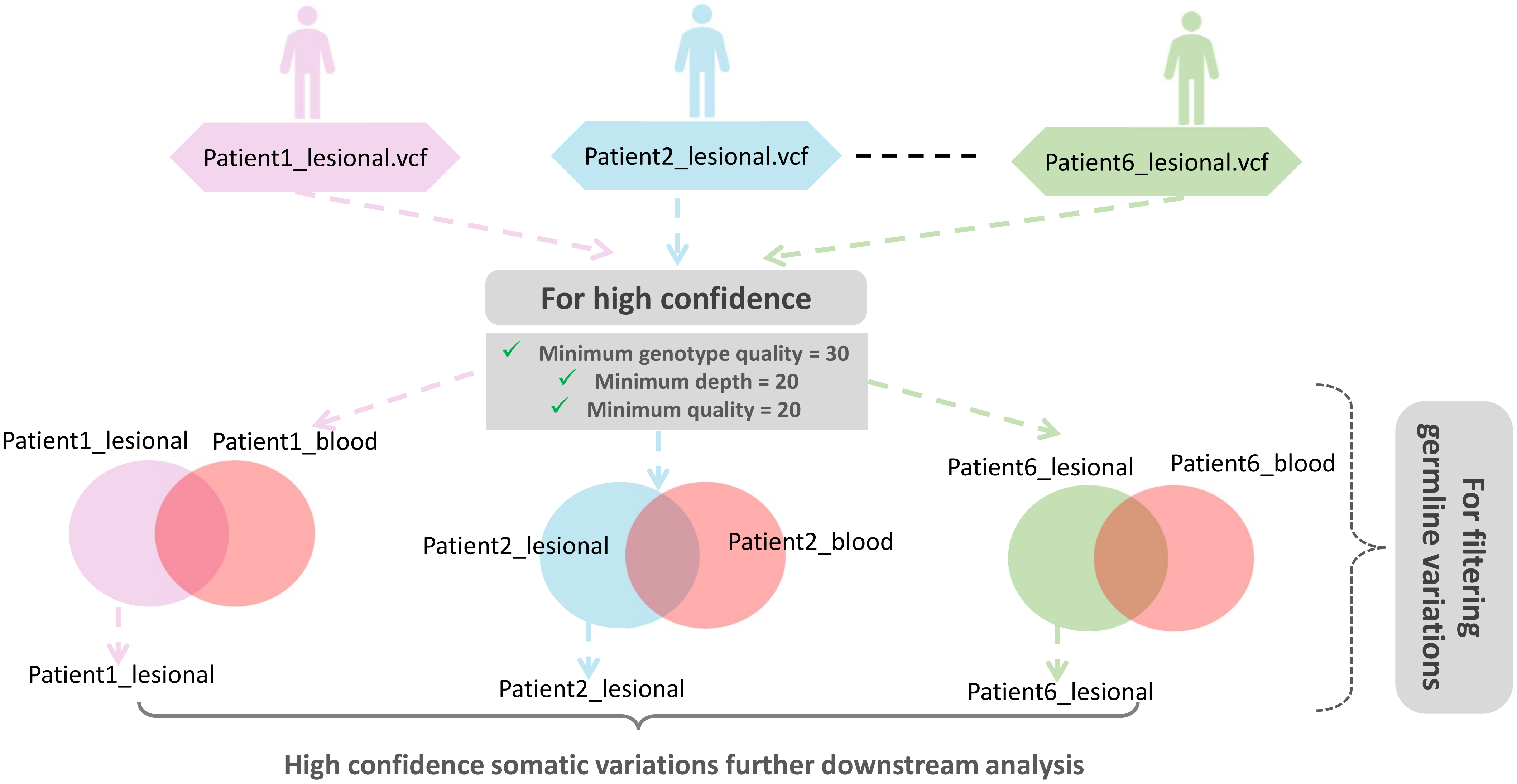


**Figure S3: Post VCF filtering steps to obtain high confidence somatic variations**

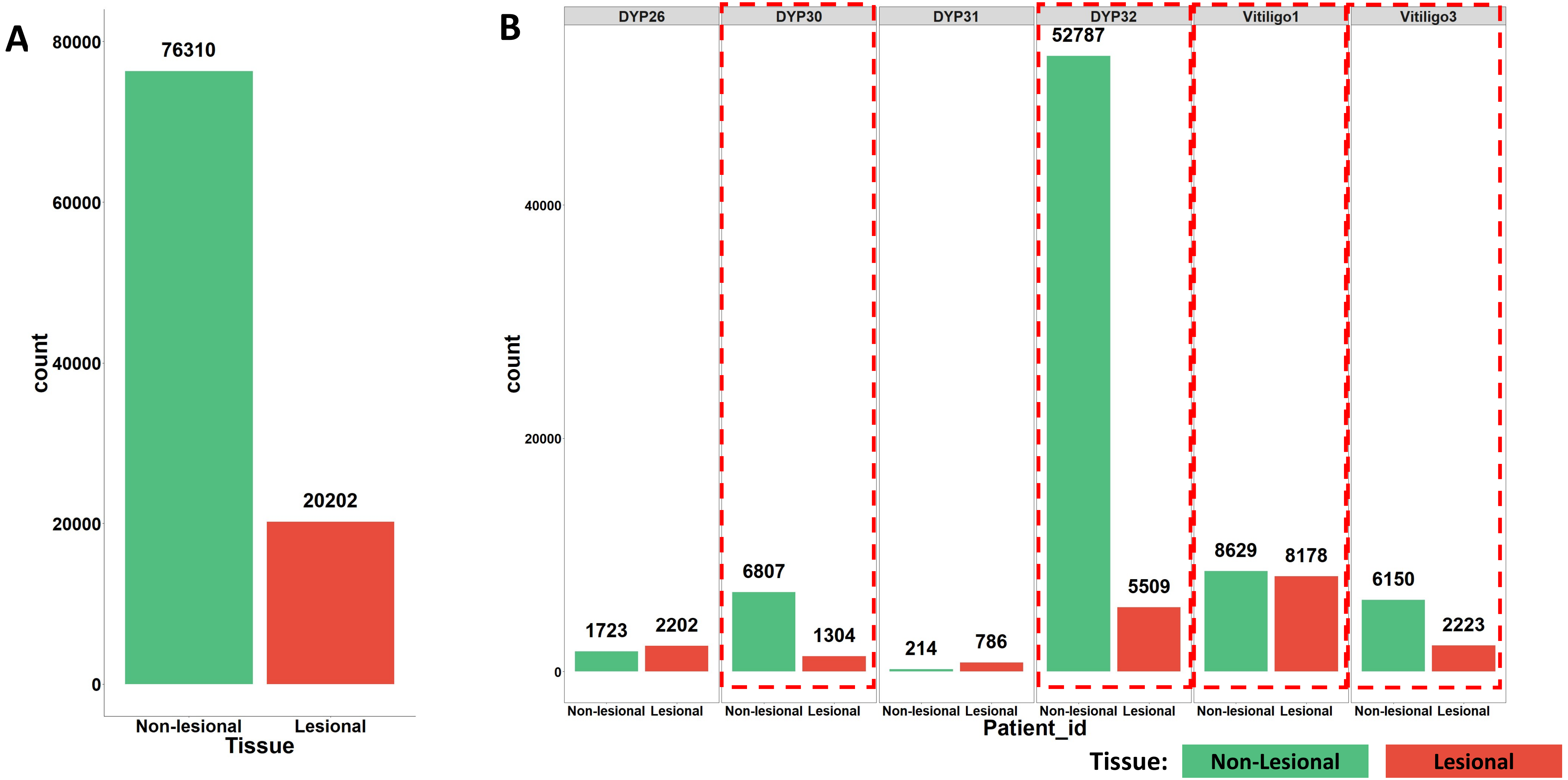


**Figure S4: Distribution of variations across lesional and non-lesional tissues showing despite non-lesional tissue being sun protected the number of variations were higher compared to the sun exposed and diseased lesional tissue (in 4 out of 6 subjects). (A)** Cumulative distribution of variations in the 2 tissues. **(B)** Sample wise distribution of variations across 6 Vitiligo subjects.

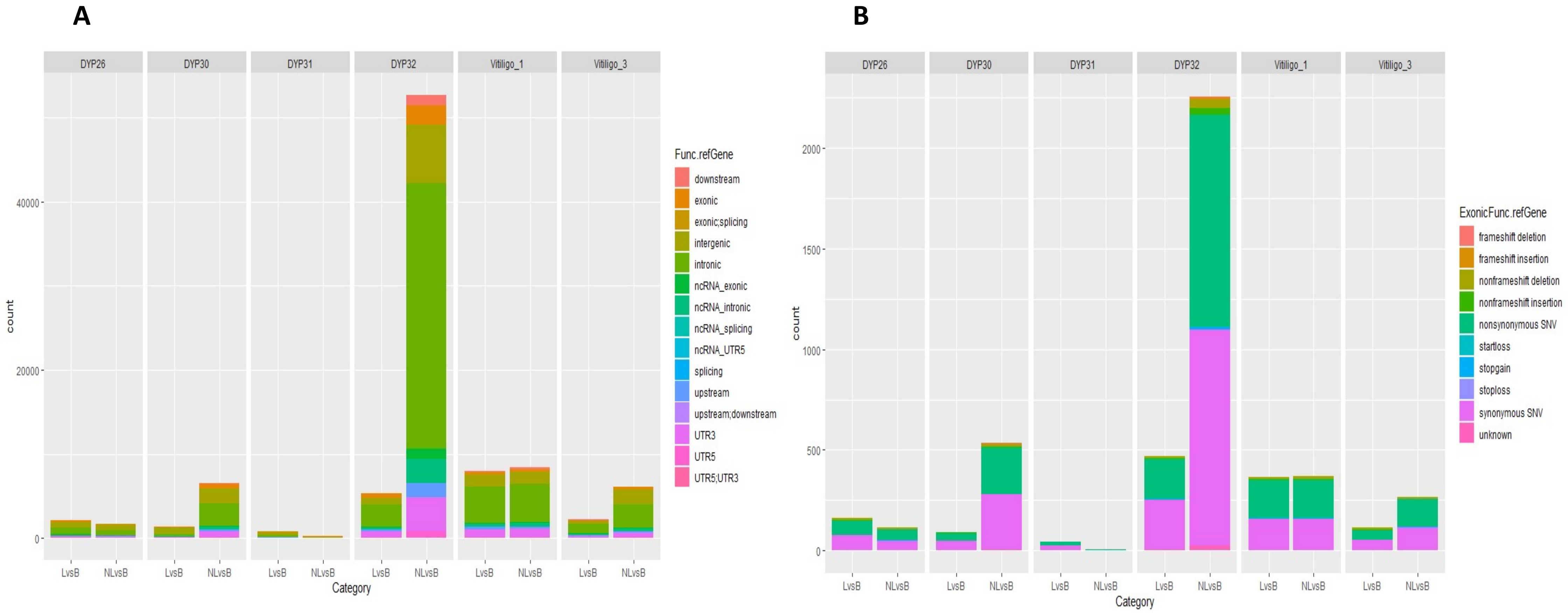


**Figure S5: Distribution of variations by class across Vitiligo samples. (A)** Variations in exonic and intronic region **(B)** Variations in exonic region

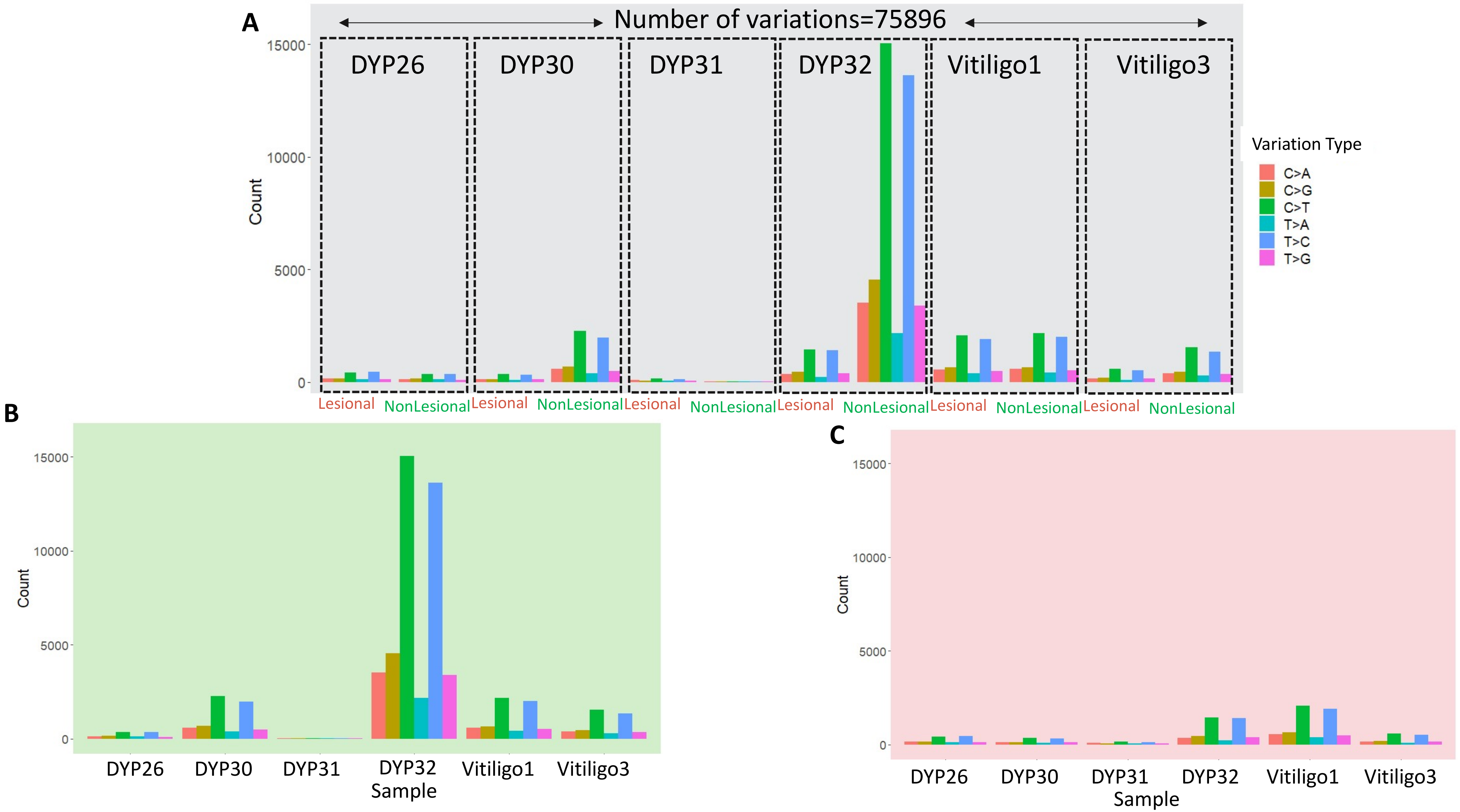


**Figure S6: Distribution of 6 variation types (A)** All samples **(B)** Non-lesional samples **(C)** Lesional samples

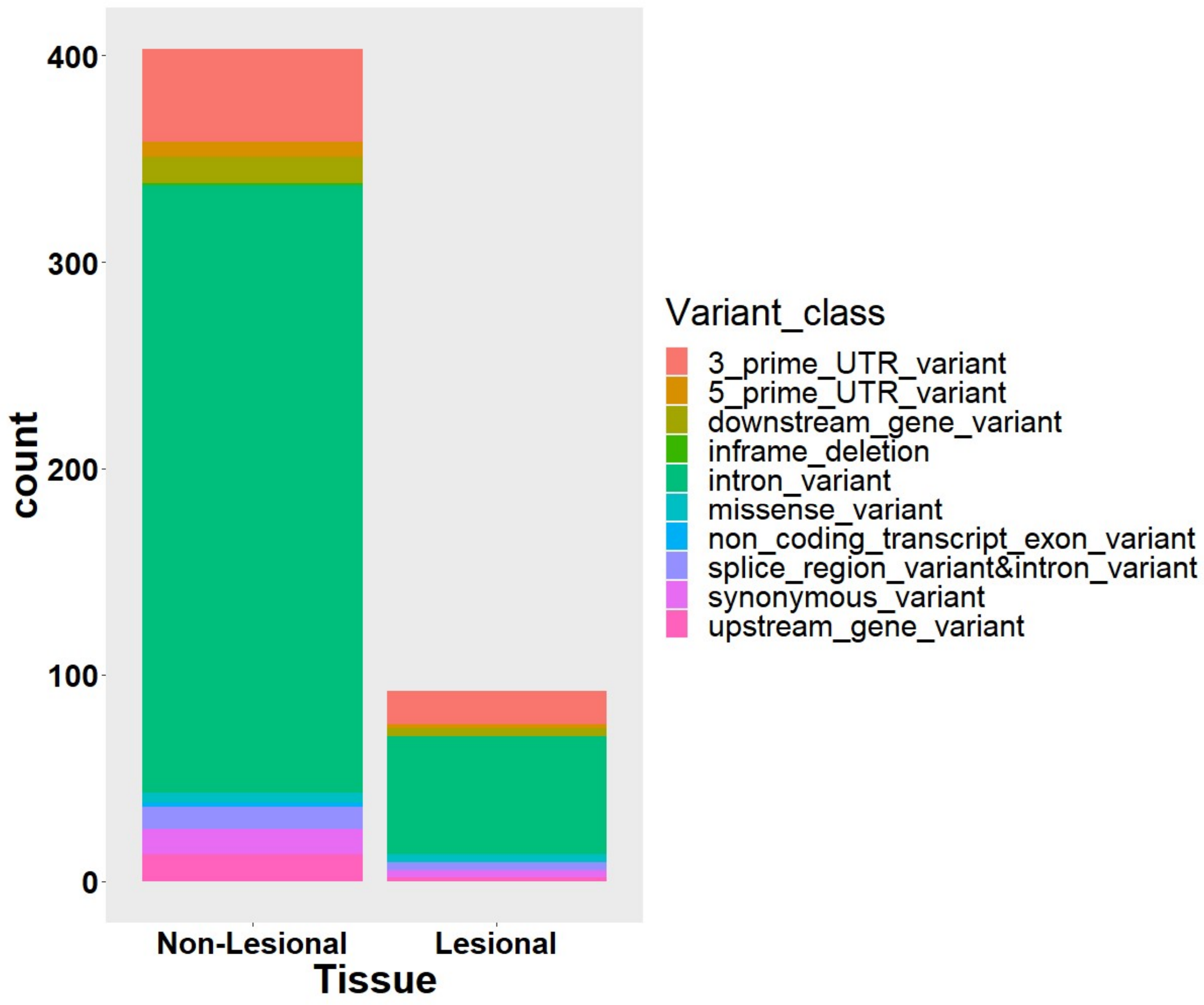


**Figure S7: Number of variations in cancer causing genes in Vitiligo non-lesional and lesional tissue (by class)**

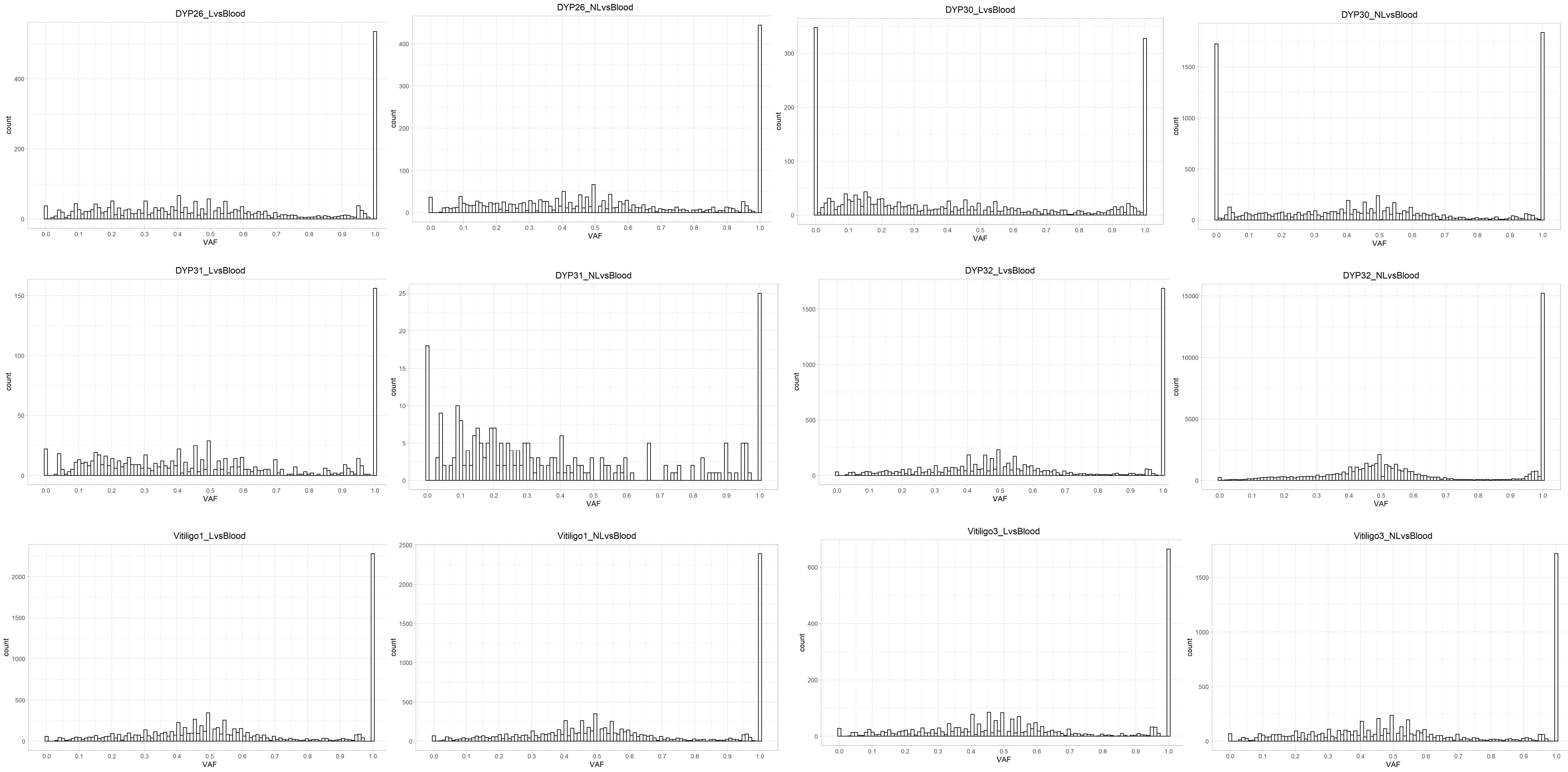


**Figure S8: Variant allele fraction(VAF) across Vitiligo samples**

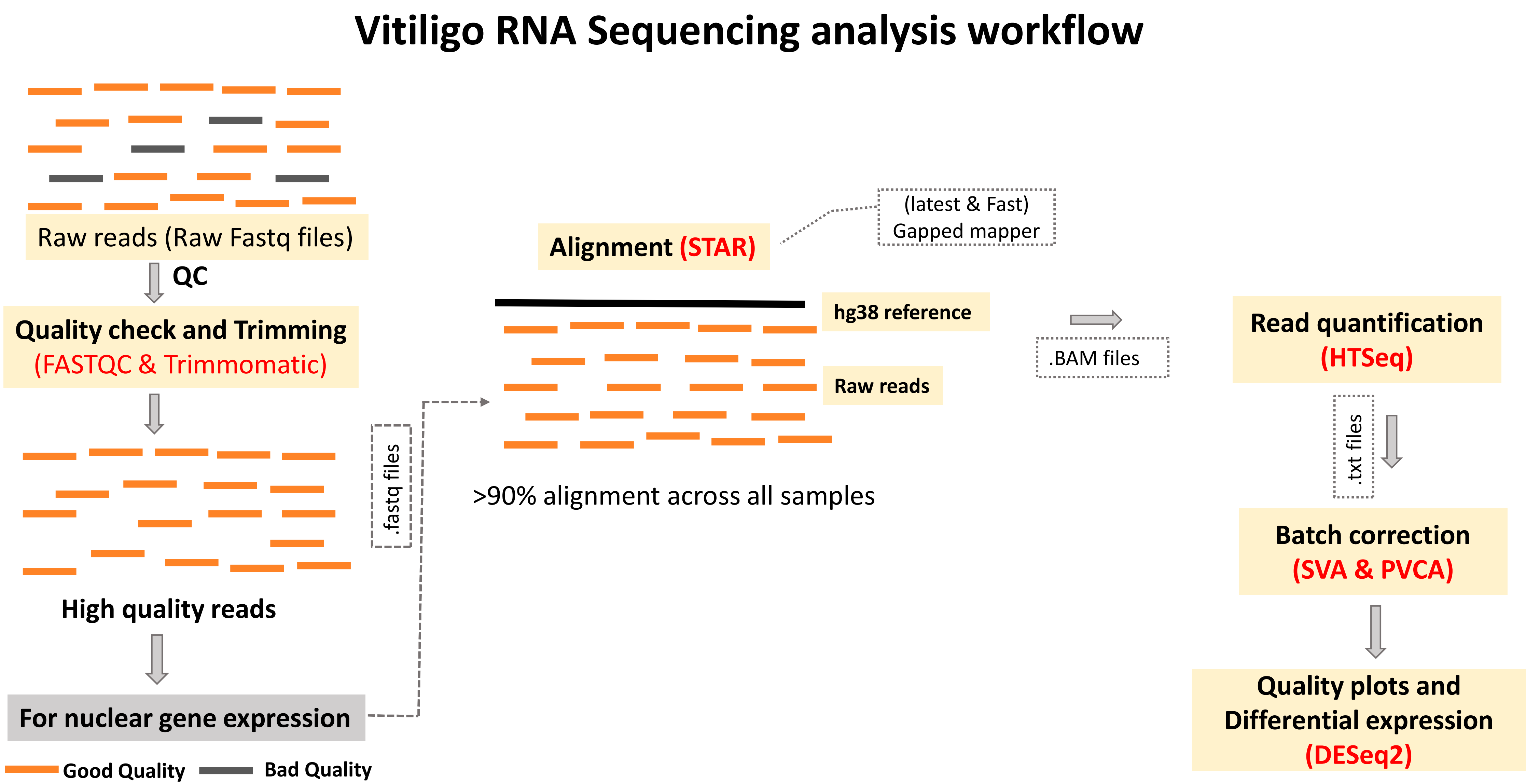


**Figure S9: Workflow followed for the transcriptome sequencing data analysis: From Fastq file till final normalized gene expression values**

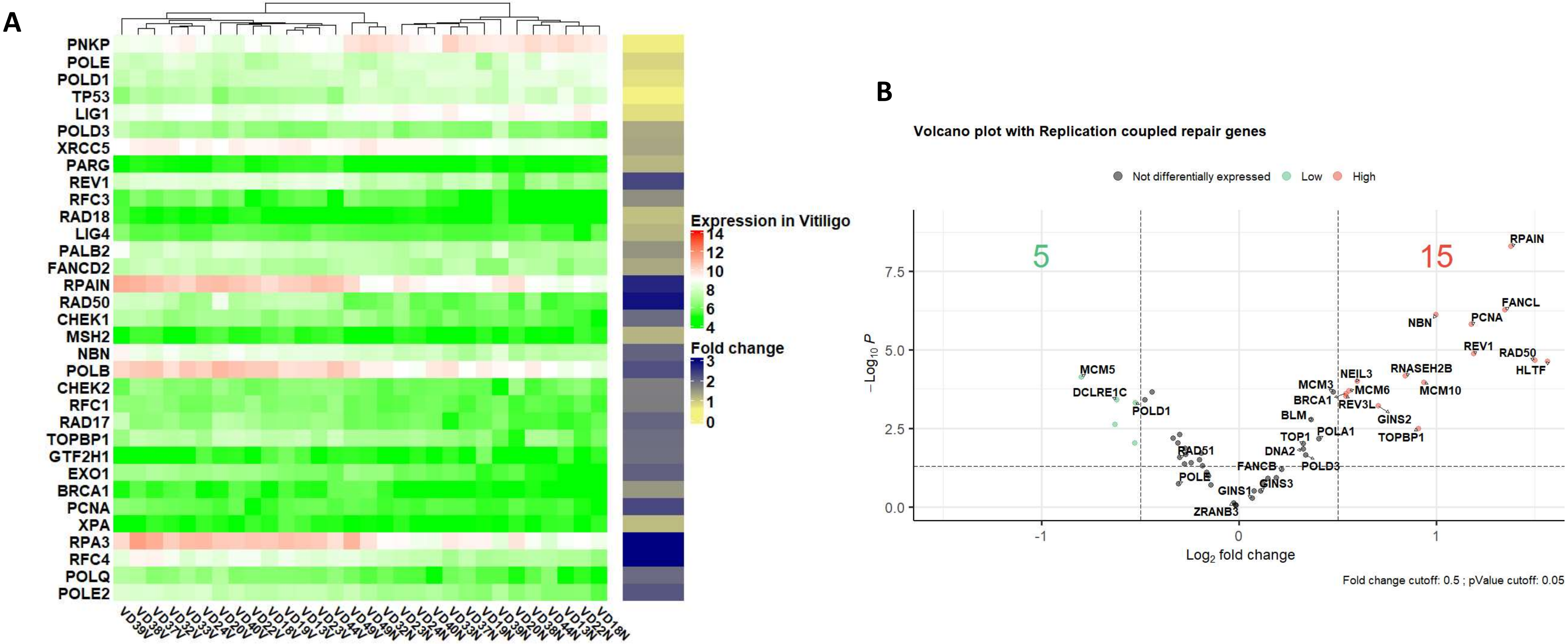


**Figure S10:Increased DNA damage & repair in Lesional tissues via replication coupled repair. (A)** Heatmap showing the expression pattern of DNA damage genes Vitiligo microarray samples. The green-red color represents the normalized expression values and the yellow-blue annotation bar is the Lesional vs Non-Lesional fold change for that gene. **(B)** Volcano plot for genes pertaining to replication coupled repair pathways in vitiligo microarray dataset. Here red dot represents a gene to be significantly up in Lesional sample whereas, green dot represents significant downregulation.

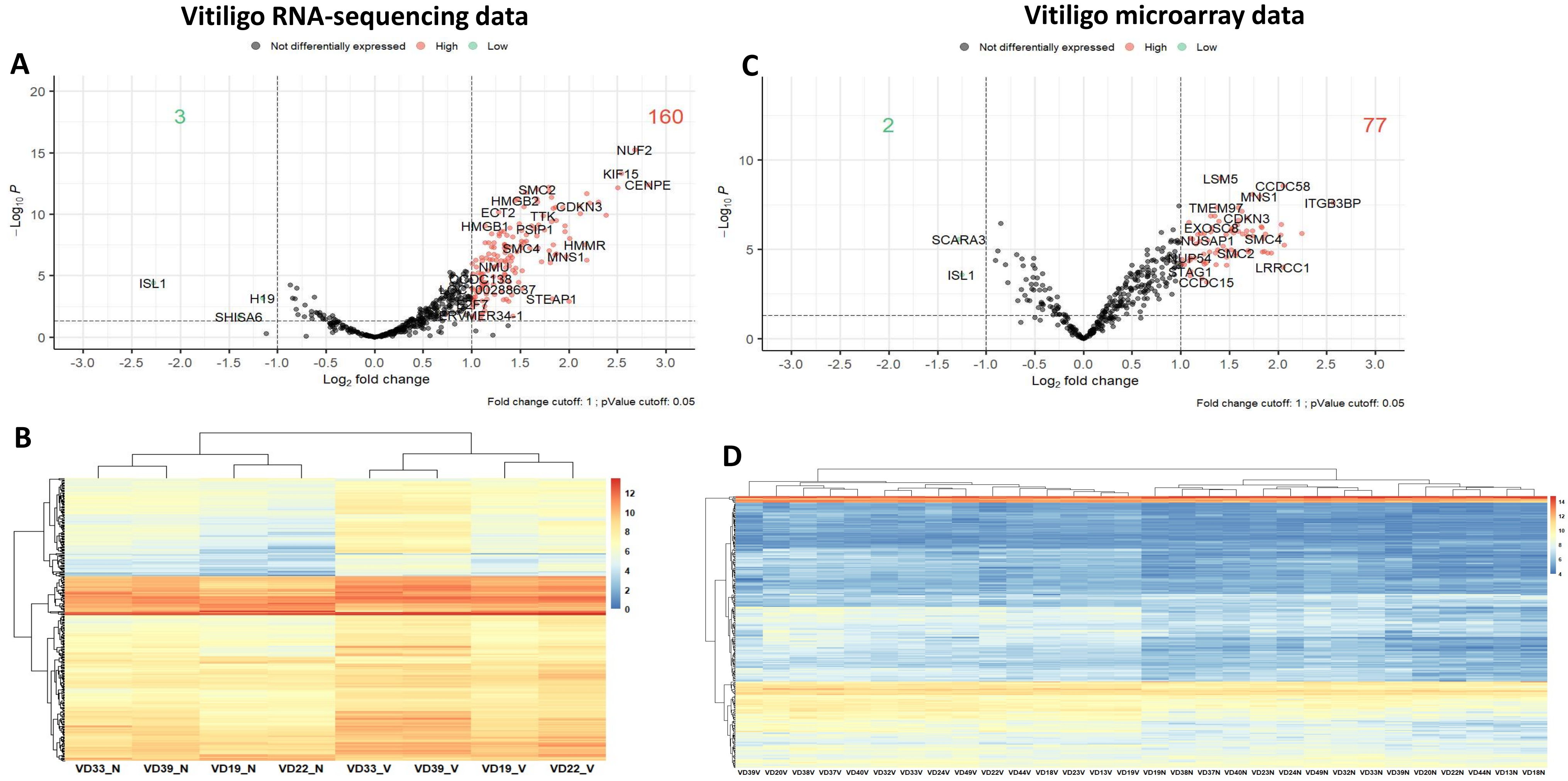


**Figure S11: Mapping of holoclone signature genes in Vitiligo datasets**. A) & B) Volcano plot showing the status of holoclone signatures in Vitiligo RNA-seq and Microarray datasets. C) & D) Heatmap showing that holoclone signatures are sufficient to segregate the Lesional and Non-Lesional tissues in RNAseq and Microarray datasets.

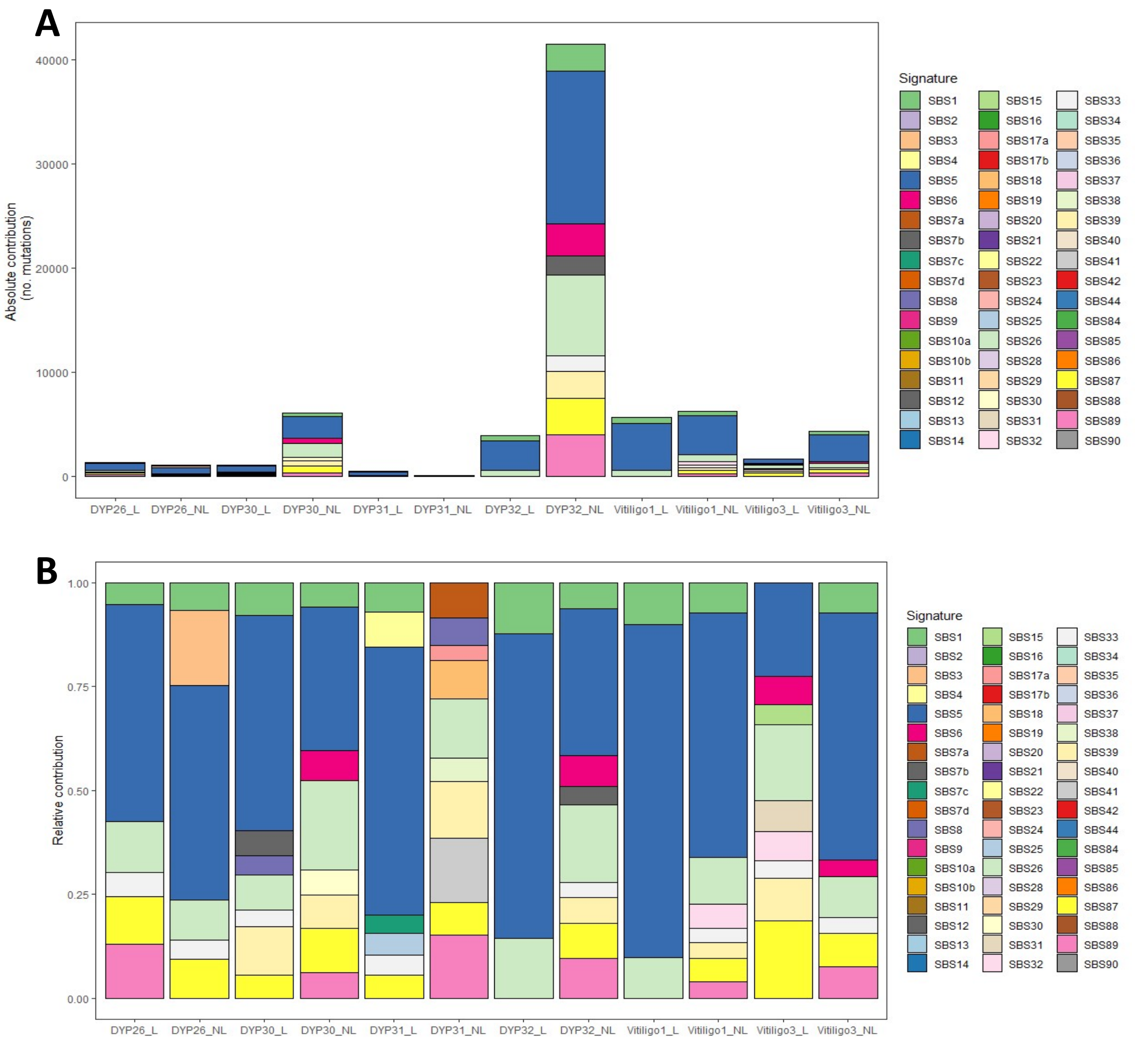


**Figure S12: Signature contribution across vitiligo samples (A)** Absolute contribution of signatures across vitiligo samples **(B)** Relative contribution of signatures across vitiligo samples
